# A Transferable Genomic Language Model Framework for Fungal Gene Essentiality Prediction

**DOI:** 10.64898/2026.09.03.749190

**Authors:** Chen Liao, Hannah G. Thomas

**Affiliations:** Department of Microbiology and Immunology, Geisel School of Medicine, Dartmouth, Hanover, New Hampshire, USA

**Author notes:** **Corresponding author** Chen Liao.

**Keywords:** gene essentiality, fungi, genomic language models, Evo2, artificial intelligence, machine learning, protein-protein interaction, drug discovery

## Abstract

Predicting biological function from genomic sequence remains a major challenge in computational and systems biology. Here, we tested whether the genomic language model Evo2, which encodes context-dependent DNA sequence patterns into embeddings, enables prediction of essential genes in fungi, a phenotype central to fungal biology and antifungal target discovery. We found that model performance was constrained not by the type or complexity of the downstream classifier, but by the biological information contained in Evo2 DNA embeddings. Specifically, the information recoverable from these embeddings progressively declined for biological features further downstream of DNA sequence, revealing a bottleneck for predicting higher-order cellular phenotypes. We alleviated this bottleneck by integrating Evo2 embeddings with two sequence-informed, system-level features: ortholog-based essentiality and protein-protein interactions. The multimodal framework demonstrated consistent performance both within and across three evolutionarily divergent yeasts (*Candida albicans*, *Saccharomyces cerevisiae*, and *Schizosaccharomyces pombe*), and its predictions were supported by published experimental evidence when transferred to the filamentous mold *Aspergillus fumigatus*. These results establish our framework as a transferrable tool for predicting essential genes across fungal genomes, including species with limited or no experimentally determined essentially data.

**Importance:** Essential genes in fungi are promising targets for antifungal drug development, yet experimental, genome-wide essentiality screening remains slow, resource intensive, and difficult to scale. As a result, comprehensive gene essentiality profiles exist for only a few model species, including *Candida albicans*, *Saccharomyces cerevisiae*, and *Schizosaccharomyces pombe*. In this work, we demonstrate that genomic language models can be leveraged through transfer learning to predict fungal essential genes directly from genomic sequence, either without training data from the target species or after fine-tuning on limited target species data. Our transfer learning approach offers a promising strategy to accelerate antifungal target discovery across understudied fungal pathogens and clinical isolates.

---

A central challenge in computational and systems biology is predicting microbial phenotypes from genomic sequences (1, 2). Although large collections of sequenced and annotated fungal genomes are now available (3), translating those sequences into accurate predictions of biological phenotypes remains difficult. This is because they may emerge from complex, multi-layered processes linking DNA variation to molecular activity, cellular physiology, and ultimately growth and morphology in specific environments (4, 5). Decoding these emergent phenotypes often benefits from integrating functional multi-omic data, including transcriptomics, proteomics, and metabolomics, with DNA sequences (6).

This challenge is best exemplified by gene essentiality, which determines whether a gene is required for viability under specific conditions (7). As a classic systems-level phenotype, essentiality can rarely be inferred from local sequence motifs alone. Instead, it emerges from the interplay of cellular networks, functional redundancy, environmental context, genetic background, and evolutionary history (8–10). While these layered dependencies make essentiality difficult to predict, identifying essential genes is vital for discovering novel antifungal drug targets and revealing how pathogens survive within human hosts (11, 12). Thus, predicting gene essentiality provides a rigorous testing ground for advancing sequence-to-function modeling in medical mycology.

Previous studies have established that machine learning can predict gene essentiality in fungi and other eukaryotes with useful accuracy (13–23). In addition to sequence-derived properties, these models frequently use orthology, evolutionary conservation, gene expression profiles, protein-protein interactions, or transposon-mutagenesis data. However, comprehensive functional genomic and experimental datasets are available for only a small number of well-studied fungal species. Models that depend on these resources can therefore be difficult to apply across species, strains, and genome assemblies, motivating approaches that predict gene essentiality directly from genomic sequence.

The success of this sequence-based strategy depends critically on how genomic DNA is represented. Because gene essentiality is an outcome of complex, high-order cellular processes, an effective representation must capture both short sequence motifs and long-range contextual relationships. Conventional DNA-based essentiality models used one-hot encoding or manually engineered sequence features (16, 24). One-hot encoding preserves nucleotide identity and order but does not by itself model dependencies between sequence positions, whereas engineered features depend on assumptions about which properties are essentiality relevant. Therefore, these representations may not fully capture the combination of regulatory, structural, and evolutionary signals associated with gene essentiality.

Recent advances in genomic language models (GLMs) offer a promising alternative by learning contextual DNA representations directly from sequence (25, 26). Pretrained on large and diverse collections of genomic sequences spanning multiple domains of life, GLMs generate embeddings, i.e., numerical representations that encode patterns learned from sequence context, without requiring manual feature engineering (**Fig. 1**). Models such as Evo2 (27) have demonstrated the utility of these learned embeddings across a range of genomic prediction tasks using human data. In this study, we evaluated the potential of Evo2-derived DNA embeddings for fungal gene essentiality prediction by systematically comparing embedding layers, sequence context lengths, and pooling strategies. All tuning of embedding generation configurations was performed in *Candida albicans* (*C. albicans*), which served as the reference species for selecting the configuration used in subsequent modeling.

**Figure 1.**
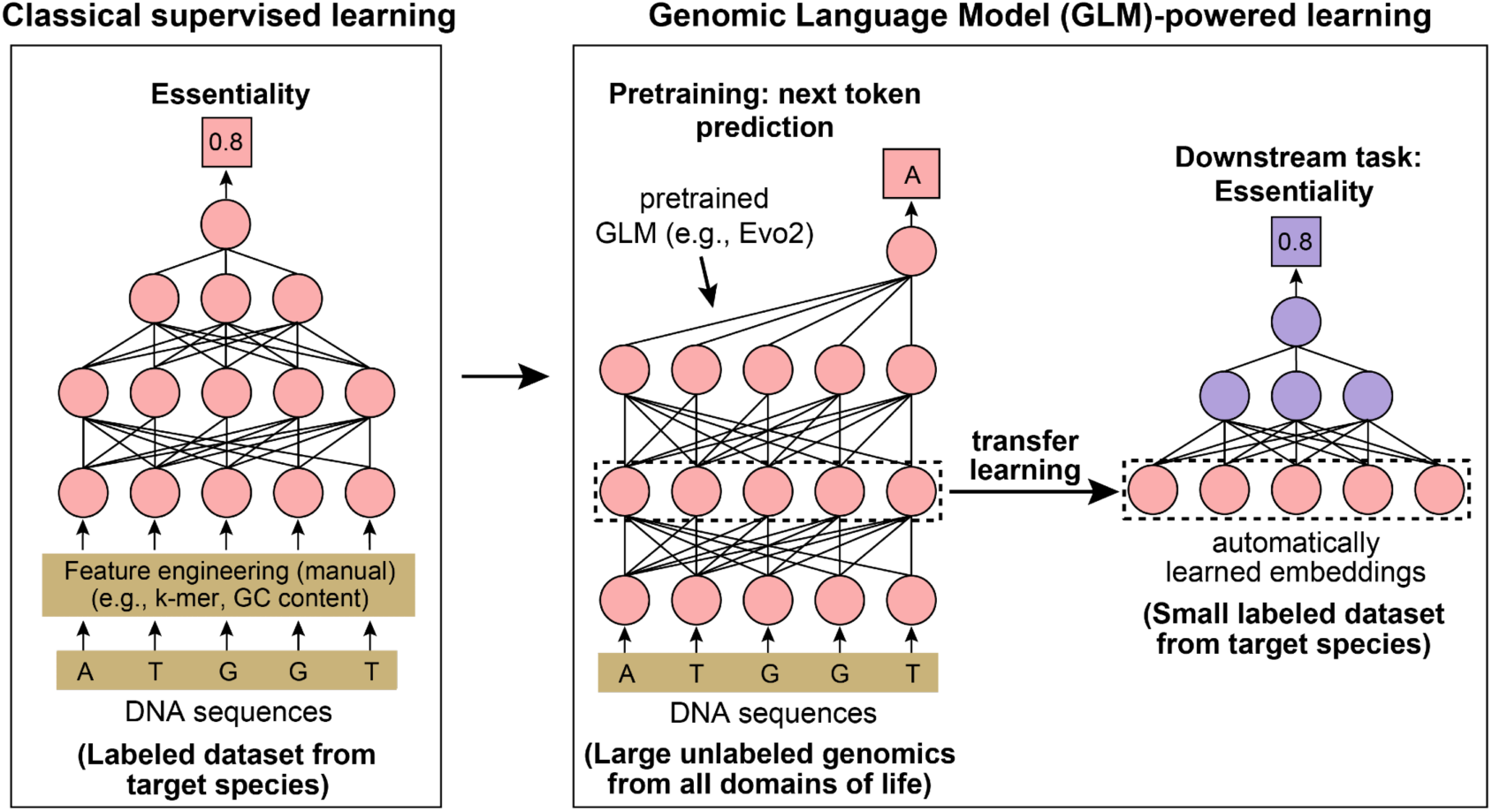
Comparison of classical and genomic language model (GLM)-powered machine learning approaches. **Left,** Classical approaches rely on one-hot encoding or feature engineering for training a supervised machine learning model. **Right,** GLM-powered approaches use a pretrained GLM to process input DNA sequences and generate sequence embeddings. These embeddings are then used as input features for downstream prediction models, which are typically smaller in size and may require less training data.

In addition to embedding generation, we developed a multimodal modeling framework that integrates the optimized embeddings with two complementary sequence-informed predictors: protein-protein interactions (PPIs) and essentiality inferred through orthology. For the ortholog-based predictor, we used genome-wide essentiality annotations for three reference fungi: *C. albicans* SC5314, *Saccharomyces cerevisiae* (*S. cerevisiae*) S288C, and *Schizosaccharomyces pombe* (*S. pombe*) 972 h- (**Table S1**). We evaluated the model independently within each reference species and assessed cross-species generalization by training on two species and predicting essentiality in the third. Our model achieved accurate and consistent performance both when no labeled data from the target species were used for training and when training included a small number of labeled genes from the target species. Finally, we combined data from all three reference species to generate genome-wide predictions of essentiality for *Aspergillus fumigatus* (*A. fumigatus*) strain A1163, a major human fungal pathogen (28) for which only a few dozen essential genes have been experimentally validated (11, 29–31).

## Results

### Evo2 identifies gene essentiality signals without supervised training

We first tested whether Evo2 could distinguish essential from non-essential genes in *C. albicans* without being trained on gene essentiality data, which is known as a “zero-shot” test. We chose the pretrained Evo2-7b-base model (a 7-billion parameter model in its base pretrained form) for this task. Evo2 was pretrained to recognize statistical patterns in DNA sequences across many organisms, enabling it to assign each sequence a likelihood score. For each *C. albicans* gene, we compared the score of the original DNA sequence with the score after introducing mutations. We tested two mutation strategies: introducing multiple premature stop codons and deleting the entire protein-coding regions. We also tested different amounts of surrounding DNA, as nearby sequences may affect how Evo2 scores a gene: the open reading frame (ORF) alone, the ORF plus nearby flanking intergenic sequence, and a larger ∼8 kb (8,192) genomic window that matches Evo2-7b-base context length (**see Methods**). A larger drop in score after mutation suggested that the gene is less able to tolerate changes, indicating it may be under stronger functional constraint and more likely to be essential.

Across all mutation strategies and sequence-context settings, AUROC (area under the receiver operating characteristic curve) values ranged from 0.597 to 0.645, and AUPRC (area under the precision-recall curve) ranged from 0.249 to 0.271 (**Table 1**). AUROC measures how well a model distinguishes essential from non-essential genes across all possible classification thresholds, with 0.5 indicating random discrimination. AUPRC measures how effectively a model identifies the essential gene class by balancing precision (the fraction of predicted essential genes that are truly essential) and recall (the fraction of true essential genes that are recovered). The random baseline for AUPRC depends on the fraction of essential genes. In this dataset where 14.7% of genes are essential, the expected AUPRC for random ranking is 0.147. Across both evaluation metrics, models incorporating the full ∼8 kb genomic context performed best, achieving higher AUROC and AUPRC values than models based on the ORF alone or the ORF plus nearby flanking intergenic sequences. Although the absolute AUPRC values are modest, they were 1.694- to 1.844-fold higher than the baseline expected from a null model that ranks genes randomly (i.e., 0.147). These results indicate that Evo2 can detect DNA sequence features associated with gene essentiality, even without being trained on essentiality labels.

**Table 1.** Evo2 performance for zero-shot gene essentiality prediction across mutation strategies and sequence contexts. Complete ORF deletion could not be evaluated in the ORF-only setting because removal of the entire ORF leaves no sequence for Evo2 scoring (marked as n/a). Normalized AUPRC was calculated as the observed AUPRC divided by the baseline AUPRC expected when genes are ranked randomly.

| Mutation Strategy | Context Window | AUROC | AUPRC | Normalized AUPRC |
| --- | --- | --- | --- | --- |
| Add stop codons | ORF only | 0.638 | 0.249 | 1.694 |
|  | ORF + intergenic | 0.618 | 0.258 | 1.755 |
|  | 8k context | 0.645 | 0.270 | 1.837 |
| Deletion | ORF only | n/a |  |  |
|  | ORF + intergenic | 0.597 | 0.263 | 1.789 |
|  | 8k context | 0.642 | 0.271 | 1.844 |

### Non-terminal Evo2 embeddings with flanking context and ORF-level mean pooling yield the best supervised classifier performance

Because Evo2 captures essentiality-related signals without task-specific training, we next tested whether its internal embeddings could support supervised gene essentiality prediction in *C. albicans*. We trained a lightweight multilayer perceptron (MLP) and used five-fold cross validation to optimize the embedding generation configuration (**Fig. 2A**). Specifically, we compared Evo2 extraction layers 20, 22, 24, 26, 28, and 30. Throughout the study, Evo2 layer numbers refer to the zero-based block indices used in the model implementation (e.g., layer 26 corresponds to blocks.26). We also compared two sequence inputs: the ORF only and an ∼8 kb genomic context window containing the ORF and its flanking sequence. Finally, we varied the pooling approach used to summarize nucleotide-level embeddings into a single gene-level representation. For the ∼8kb context, we compared pooling over the ORF only, the ORF plus intergenic regions, or the entire ∼8 kb window, each using mean pooling, max pooling, or final-token selection. For the ORF-only input, the pooling region comprised the entire ORF, and the same three methods were evaluated.

**Figure 2.**
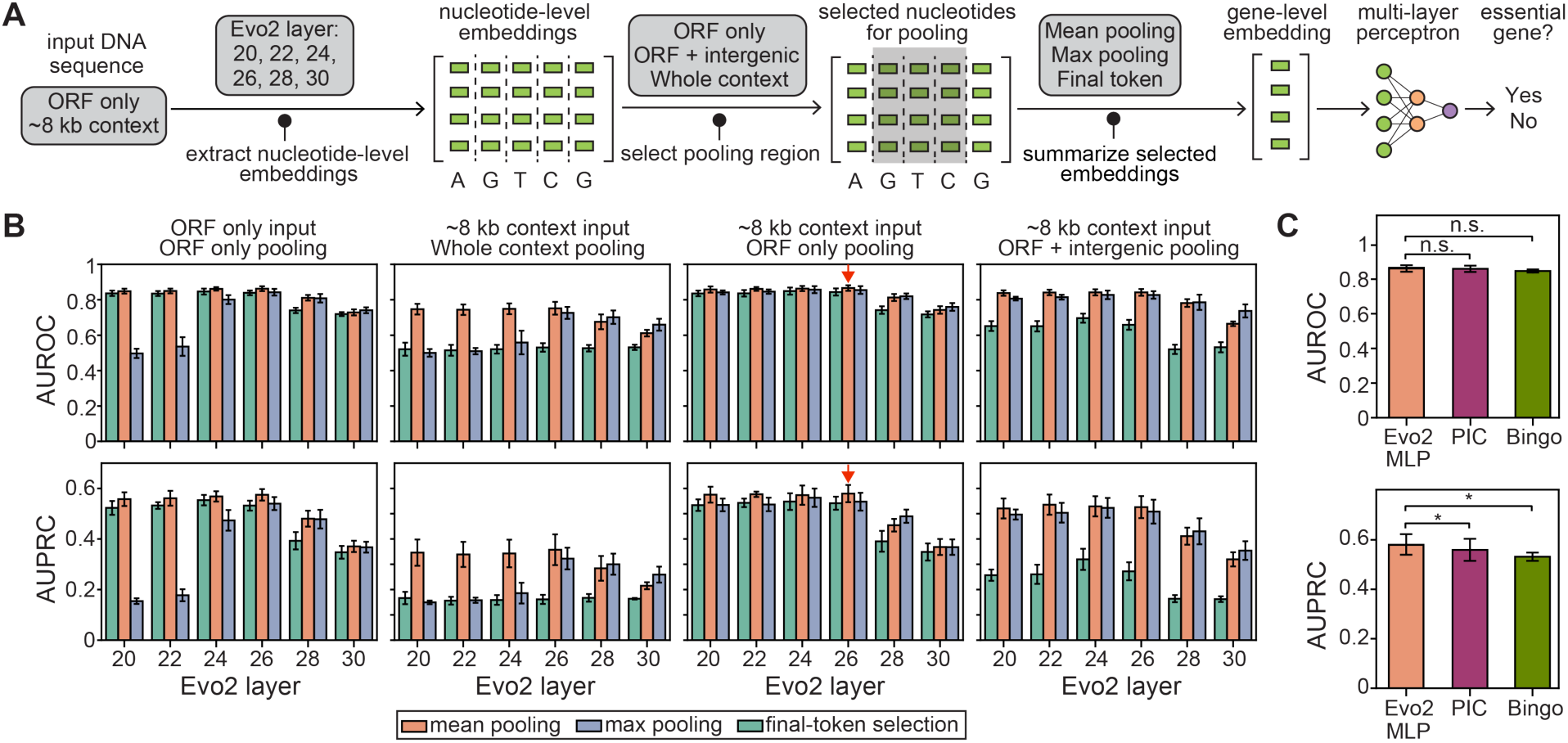
Optimization and benchmarking of supervised gene essentiality prediction using Evo2 embeddings and a lightweight MLP. **A**, Overview of the embedding generation and essentiality classification workflow. Gene-level representations were generated from frozen Evo2 embeddings by systematically varying the input sequence context, embedding extraction layer, nucleotide pooling region, and summarization method. These representations were then used to train an MLP classifier on labeled essentiality data. **B**, MLP performance (upper row: AUROC; lower row: AUPRC) trained with different embedding generation configurations evaluated by five-fold cross validation. Red arrows indicate the configuration with the highest mean AUROC and AUPRC. **C**, Comparison of the best-performing Evo2-MLP configuration with PIC and Bingo, under the same five-fold cross validation setting. Bars show the mean across folds, and error bars indicate standard deviation. n.s., not significant; *, P < 0.05 (paired *t*-test).

The configuration with the highest AUROC and AUPRC used embeddings from Evo2 layer 26, sequences with ∼8 kb of surrounding context, and mean pooling over the ORF region (**Fig. 2B**; **Table S2**). Performance declined in the final Evo2 layers, likely because these embeddings are more specialized for Evo2’s pretraining task of autoregressive next-token prediction (i.e., predicting each next nucleotide from the preceding DNA sequence) and are less suitable for downstream essentiality prediction. Notably, layer 26 was also identified as effective for exon classification in the original Evo2 study, suggesting that it contains sequence features that transfer across downstream tasks. Including ∼8 kb of context around each ORF also produced more consistent performance across pooling methods (**Fig. 2B**, third column). However, pooling over the broader flanking sequence reduced performance, likely because non-coding regions introduce noise into the gene-level representations (**Fig. 2B**, second column). Although pooling was restricted to the ORF, the surrounding context could still affect the embeddings of nucleotides within the ORF and therefore the final gene-level representation.

Using the best-performing configuration, we compared our Evo2-based model with PIC (32) and Bingo (33), two related sequence-based gene essentiality prediction models that use protein language model representations (**Table S3**). Across five-fold cross validation, our Evo2-based model achieved the highest mean AUROC and AUPRC. While the AUROC differences were modest (mean AUROC: 0.864 for Evo2, 0.862 for PIC, and 0.850 for Bingo), the improvement was more pronounced and statistically significant for AUPRC (mean AUPRC: 0.582 for Evo2, 0.560 for PIC, and 0.533 for Bingo), corresponding to relative improvements of 3.9% over PIC and 9.2% over Bingo. For reference, the expected AUPRC for random ranking in this dataset is 0.147, such that all three models achieved AUPRC values more than 3.5-fold above random. Notably, the Evo2-based model used only a lightweight MLP classifier on frozen Evo2 embeddings, whereas PIC and Bingo use more specialized downstream architectures, including multi-head attention in PIC and a graph neural network in Bingo. Together, these results show that training on essentiality labels substantially improved prediction compared to direct Evo2 sequence scoring, increasing the maximum mean AUROC from 0.645 to 0.864 and the maximum mean AUPRC from 0.271 to 0.582. Achieving this performance depends strongly on how the embeddings were generated. With the optimized embedding strategy, frozen Evo2 embeddings paired with a lightweight MLP matched or modestly outperformed two protein language model-based benchmarks, indicating that competitive performance does not require a specialized, sophisticated downstream classifier architecture.

### Increasing MLP complexity or changing classifier type does not improve supervised performance

After showing that a lightweight MLP was sufficient to extract predictive information from Evo2 embeddings, we moved on to explore whether increasing classifier capacity, by enlarging the hidden layers and thereby increasing the number of trainable parameters, could yield further improvements. We trained MLPs with progressively larger hidden layers using the same frozen Evo2 DNA embeddings to predict essential genes in *C. albicans* and evaluated mean AUROC and AUPRC across five-fold cross validation (**Tables 2**). Performance remained largely unchanged as the first hidden layer increased from 64 to 4,096 neurons and the second hidden layer increased from 8 to 256 neurons. Across these changes, mean AUROC varied only from 0.864 to 0.867 and mean AUPRC from 0.574 to 0.581.

**Table 2.** Effect of downstream classifier type and complexity on supervised gene essentiality prediction from Evo2 embeddings. AUROC and AUPRC are reported as mean values across the five cross validation folds. All classifiers were trained and evaluated using the same Evo2 embeddings and cross validation splits as in **Fig. 2**. The first four rows compare MLP architectures with increasing hidden layer sizes (dim1 and dim2 denote the number of neurons in the first and second hidden layers, respectively). The MLP with dim1=256 and dim2 =16 is the default configuration used in **Fig. 2**. The remaining rows compare logistic regression, k-nearest neighbors, random forest, and XGBoost, For random forest and XGBoost, performance is reported for both default and grid-search-optimized hyperparameter settings.

| Model | Hyperparameters | Mean AUROC | Mean AUPRC |
| --- | --- | --- | --- |
| Multi-layer perceptron (MLP) | dim1=64, dim2=8 | 0.864 | 0.574 |
|  | dim1=256, dim2=16<br>(configuration used in **Fig. 2**) | 0.865 | 0.581 |
|  | dim1=1,024, dim2=64 | 0.864 | 0.578 |
|  | dim1=4,096, dim2=256 | 0.867 | 0.576 |
| Logistic regression | default | 0.862 | 0.559 |
| K-nearest neighbor | default | 0.786 | 0.421 |
| Random forest | default | 0.824 | 0.525 |
|  | optimized | 0.851 | 0.553 |
| XGBoost (eXtreme Gradient Boosting) | default | 0.851 | 0.548 |
|  | optimized | 0.863 | 0.572 |

We then compared MLP with alternative classifiers trained on the same Evo2 embeddings, including logistic regression, two tree-based models, and k-nearest neighbors. Logistic regression achieved a mean AUROC of 0.862 and a mean AUPRC of 0.559, closely matching the MLPs in AUROC but performing modestly worse in AUPRC. This similarity suggests that the MLP extracts little additional predictive information beyond that captured by a linear classifier. Two tree-based models, Random forest and XGBoost (eXtreme Gradient Boosting), also underperformed the MLPs using default hyperparameters, whereas their hyperparameter-optimized versions reached comparable performance. In contrast, k-nearest neighbors performed substantially worse (mean AUROC: 0.786; mean AUPRC: 0.421), suggesting that local proximity in the Evo2 embedding space does not reliably reflect shared essentiality labels. Together, these results show that neither increasing MLP complexity nor substituting common alternative classifiers produced meaningful performance gains. The observed performance ceiling is therefore constrained by the information encoded in the Evo2 embeddings rather than by downstream classifier capacity or model type.

### Biological features distal to DNA sequence are less recoverable from Evo2 embeddings

Evo2 embeddings encode numerical representations of DNA sequence and its surrounding context, but they may not capture all biological features related to gene essentiality. Because the information contained in these embeddings constrains model performance, we then examined which essentiality-related features are poorly represented. A previous study predicted *C. albicans* gene essentiality using a random forest model trained on 13 curated features spanning sequence composition, transcriptomic and transposon-insertion statistics, orthology to *Saccharomyces cerevisiae*, and population genetic summaries (14). To directly compare these curated features with Evo2 embeddings, we trained the same MLP classifier used in our best-performing Evo2-based model on the 13 curated features and evaluated performance using the same five-fold cross validation framework (**see Methods**). Interestingly, the curated-feature model outperformed our Evo2-based model (mean AUROC: 0.921 vs. 0.864; mean AUPRC: 0.676 vs. 0.582). Because the classifier and evaluation procedure were held constant, this comparison suggests that the 13 curated features contain essentiality-associated information that is not fully represented in Evo2 embeddings.

To determine how well each curated feature could be recovered from Evo2 embeddings, we trained a separate MLP to predict each feature and evaluated performance using the mean Spearman correlation (ρ) between predicted and observed values across five held-out test folds (**see Methods**). The 13 features showed a clear gradient of recoverability (**Table 3**). Features most directly derived from coding sequence, including coding sequence length and codon adaptation index, were predicted most accurately (mean ρ > 0.90). Features related to gene expression and transposon insertion patterns showed intermediate recoverability (mean ρ ≈ 0.60-0.66), suggesting that embeddings capture some, but not all, of the relevant information in the local DNA sequence. Features reflecting downstream biological context, including co-expression, orthology-based essentiality, synthetic sick/lethal paralog relationships, and finer-scale transposon insertion context, were recovered less accurately (mean ρ ≈ 0.35-0.53). As expected, population-level sequence variation and gene expression variance are the least recoverable features (mean ρ < 0.31), because they depend on strain-specific, environmental, or population-level information that cannot be reliably inferred from the *C. albicans* SC5314 reference genome. Together, these results show that Evo2 embeddings most effectively capture features closely linked to the underlying DNA sequence, whereas features reflecting progressively more distal biological processes and broader biological context are less recoverable.

**Table 3.** Recoverability of 13 curated essentiality-associated features from Evo2 embeddings. For each feature obtained from Fu *et al.* (14), we trained a separate MLP regression model to predict its value from Evo2 embeddings. Spearman correlation (ρ) between predicted and observed values was calculated separately for each held-out test fold, and recoverability is reported as the mean ρ across the five folds. Features are grouped into four categories according to their recoverability and biological relationship to DNA sequence. CAI, codon adaptation index; TPM, transcripts per million.

| Category | Curated Feature | Definition | Mean Spearman $\rho$ |
| --- | --- | --- | --- |
| Sequence and composition | Length | Total length of intron-free coding sequence (bp) | 0.961 |
|  | CAI | Codon adaptation index | 0.909 |
| Genomic architecture and gene expression | Gene expression median | Median gene expression (TPM) | 0.662 |
|  | Hits | Number of transposon insertion sites within the ORF | 0.619 |
|  | Freedom index | (Longest insertion-free interval within ORF) / (ORF length) | 0.599 |
|  | Neighborhood index | Number of hits per ORF normalized by the hits in surrounding intergenic sequences | 0.596 |
| Network-level interactions and systems biology | Co-expression degree | Number of co-expression partners | 0.525 |
|  | Reads | Number of sequencing reads mapping within the ORF | 0.459 |
|  | Upstream hits 100 | Number of transposon insertion sites within 100 bp upstream of the start codon | 0.380 |
|  | Synthetic sick/lethal paralogs in *S. cerevisiae* | Whether *S. cerevisiae* ortholog is duplicated and sister paralogs show synthetic sick/lethal interactions | 0.356 |
|  | Ortholog essentiality in *S. cerevisiae* | Whether *S. cerevisiae* ortholog is essential | 0.349 |
| Environment and population genetics | Sequence variation | SNPs per nucleotide (normalized) | 0.302 |
|  | Gene expression variance | Variance of gene expression (TPM) | 0.188 |

### Ortholog essentiality and predicted PPI networks provide complementary inputs to Evo2 embeddings

Since Evo2 embeddings may miss high-order biological information, improving essentiality prediction may require complementary features that capture signals not represented in these embeddings. We reasoned that the most informative complementary features would be poorly recoverable from the embeddings yet strongly predictive of gene essentiality. To identify such features, we performed a leave-one-feature-out analysis of the complete 13-feature model and quantified changes in AUROC and AUPRC after removing each feature individually (**Fig. S1**; **see Methods**). *Saccharomyces cerevisiae* ortholog essentiality best met both criteria: it was weakly recoverable from Evo2 embeddings (mean ρ = 0.349), yet its removal reduced AUROC by 0.008 and AUPRC by 0.035 (biggest drops among all features for both metrics). Because orthologous relationships between *C. albicans* and *S. cerevisiae* can be computationally inferred from their genome sequences, this feature can be incorporated without requiring additional experimental data from *C. albicans*.

Protein-protein interaction (PPI) networks may also provide complementary biological context not captured by Evo2 embeddings, including protein connectivity, local neighborhood structure, and interaction partner relationships (34). These network properties can provide information about a gene’s functional role, redundancy with other genes, and participation in essential cellular pathways, all of which may influence gene essentiality. Because our study aims to predict essentiality from DNA sequence alone, incorporating experimental PPI data falls outside our scope. However, computational methods for predicting PPs from sequence have improved substantially in accuracy, scalability, and runtime efficiency (35). We therefore used FlashPPI (36), a recently published tool for rapid inference of microbial protein interactions, to generate a predicted PPI network for *C. albicans* (**Table S4;** contact score ≥ 0.5). Comparison with STRING (37) v12.0 physical interactions at a confidence score ≥ 700, FlashPPI achieved a precision of 0.239 (23.9% of its predicted interactions were supported by STRING) and a recall of 0.078 (it recovered 7.8% of the interactions reported in STRING). Despite its limited recall, the continued progress of artificial intelligence (AI)-based PPI prediction and the potential of PPI networks to capture biological context beyond DNA sequence motivated us to include FlashPPI-predicted PPI information as an additional input.

### Multimodal integration of Evo2 embeddings, ortholog essentiality, and predicted PPI network features improves gene essentiality prediction

Based on the above analyses, we combined Evo2 embeddings with *S. cerevisiae* ortholog essentiality and predicted PPI network features into a multimodal model. Each modality was first projected into a modality-specific latent space and then combined using bilinear interaction layers to capture cross-modal dependencies before final classification (**Fig. 3A**). Orthologs between *C. albicans* and *S. cerevisiae* were inferred using OrthoFinder (38), and the predicted PPI network was represented using node2vec (39) embeddings generated from a network weighted by FlashPPI contact scores (**see Methods**). We evaluated the contribution of each modality through systematic ablation analysis, in which one or more inputs were removed while keeping the remaining architecture unchanged (**Fig. 3B; Table S5**).

**Figure 3.**
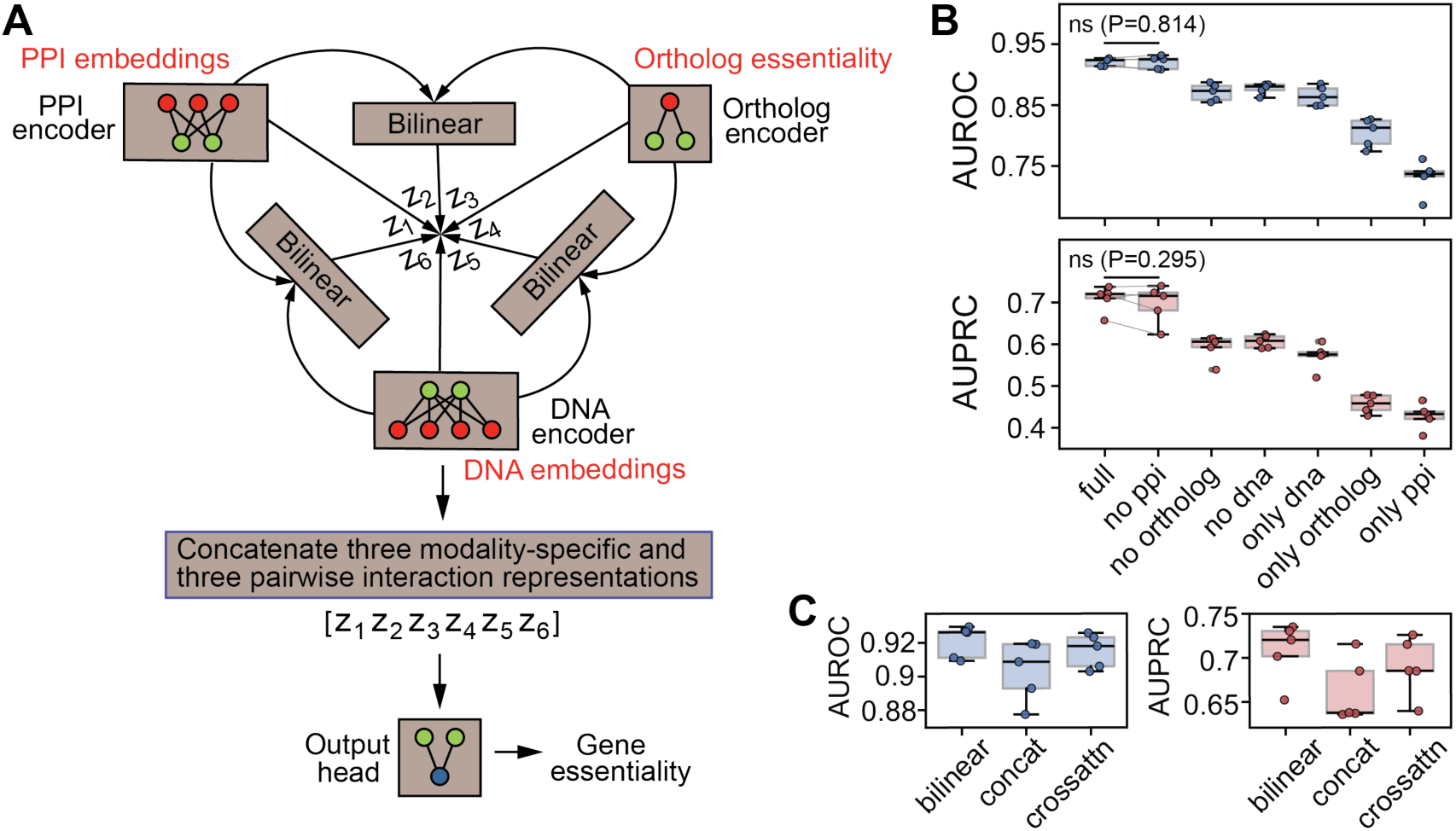
Multimodal model architecture and ablation analysis for gene essentiality prediction. **A,** Schematic of the multimodal model. Evo2 DNA embeddings, *S. cerevisiae* ortholog essentiality, and PPI-derived node2vec embeddings are projected into modality-specific latent spaces and then integrated using bilinear interaction layers. z1-z6 denote latent representations of individual modalities and their pairwise interactions. **B,** Ablation analysis showing cross-validated AUROC and AUPRC after removing one or more input modalities. “Full” denotes the model using all three modalities. “no ppi,” “no ortholog,” and “no dna” denote models with the indicated modality removed. “only dna,” “only ortholog,” and “only ppi” denote single-modality models. Each dot represents one test fold. P values were calculated using paired *t*-tests across matched cross validation folds. ns, not significant. **C,** Comparison of bilinear fusion with alternative modality fusion strategies. “Concat” denotes simple concatenation of modality-specific latent representations without explicit pairwise interactions. “Crossattn” denotes a cross-attention architecture that models dependencies among modalities. Boxplots: center lines represent medians, boxes represent the 25^th^-75^th^ percentiles, and whiskers extend to the most extreme values within 1.5 times interquartile range.

The full multimodal model achieved the best performance across five-fold cross validation (mean AUROC: 0.921; mean AUPRC: 0.709). Removing either Evo2 embeddings (mean AUROC: 0.877; mean AUPRC: 0.606) or *S. cerevisiae* ortholog essentiality (mean AUROC: 0.871; mean AUPRC: 0.593) substantially reduced performance, whereas removing PPI network information had little effect (mean AUROC: 0.920; mean AUPRC: 0.700). Single-modality models showed further reductions in performance, with the PPI-only model performing worst (mean AUROC: 0.732; mean AUPRC: 0.428). These results indicate that Evo2 embeddings provide the strongest predictive signal, while *S. cerevisiae* ortholog essentiality contributes complementary information beyond that captured by DNA sequence. Finally, bilinear fusion outperformed two alternative approaches for integrating modalities (**see Methods**): simple concatenation (mean AUROC: 0.904; mean AUPRC: 0.662) and cross attention fusion (40) (mean AUROC: 0.915; mean AUPRC: 0.690), supporting its use as the final multimodal integration strategy (**Fig. 3C**).

### The multimodal model generalizes to *S. cerevisiae* and *S. pombe*

To assess whether the multimodal model generalizes beyond *C. albicans*, we trained separate models for *S. cerevisiae* S288C and *S. pombe* 972h- and evaluated them using the same five-fold cross validation framework. For each target species, the model incorporated Evo2 embeddings, PPI-network embeddings, and ortholog-essentiality information from *C. albicans* and the remaining fungal species. The models achieved mean AUROCs of 0.889 for *S. cerevisiae* and 0.856 for *S. pombe*, with mean AUPRCs of 0.751 and 0.738, respectively. We also evaluated the contribution of FlashPPI embeddings by comparing these models with otherwise identical models lacking the interaction network features (**Fig. S2**). FlashPPI embeddings did not improve performance in *C. albicans* or *S. pombe*, but significantly increased AUPRC, although not AUROC, in *S. cerevisiae*. These results demonstrate that our modeling strategy generalizes across the three reference species and the contribution of predicted PPI network information is species-dependent.

### Nucleotide-level attribution reveals species-biased coding motifs associated with fungal gene essentiality

The ability of the multimodal models to generalize across fungal species raised the question of whether their DNA representations reflect conserved sequence motifs or species-specific coding patterns. To identify motifs associated with gene essentiality, we applied integrated gradients (41) to assign nucleotide-level contribution scores to each gene sequence, followed by motif discovery using TF-MoDISco (42) (**see Methods**). Species-specific analyses identified 12, 14, and 13 motifs of 50 nucleotides in *C. albicans*, *S. cerevisiae*, and *S. pombe*, respectively (**Table S6**). We then pooled sequences from all three species and repeated the analysis using the same pipeline to identify potentially shared fungal motifs. This analysis recovered 18 pooled motifs (**Table S6**), but most were strongly biased toward one species rather than evenly represented across all three. Of the 18 motifs, 13 were dominated by *C. albicans* seqlets and 5 by *S. cerevisiae* seqlets, with a mean dominant-species seqlet fraction of 82.40%. These results support predominantly species-biased coding sequence attribution patterns rather than a single conserved fungal motif shared across species.

Among the pooled motifs, Motif 3 showed the strongest overall association with gene essentiality (Odds Ratio = 4.15, P = 2.3e-08; two-sided Fisher’s exact test): 49.3% of motif-bearing genes are essential, compared to 19.1% of all labeled genes across the three species (**Table S6**). Although 71.9% of its seqlets originates from *C. albicans*, Motif 3 was the only pooled motif whose motif-bearing genes includes an orthologous group represented by one gene from each of the three species (**see Methods**). The orthologous group corresponds to RPC10, which encodes the conserved ABC10-alpha subunit shared by RNA polymerases I, II, and III. **Fig. 4A** shows the nucleotide-level attribution patterns of Motif 3 within RPC10 orthologs from each species. The attribution patterns contained nucleotides with positive or negative contributions to the predicted essentiality and showed an apparent three-nucleotide periodicity, reflecting the structure of the coding frame (**Fig. 4B**). Other than Motif 3, we did not identify broadly supported cross-species motifs associated with conserved orthologous gene families.

**Figure 4.**
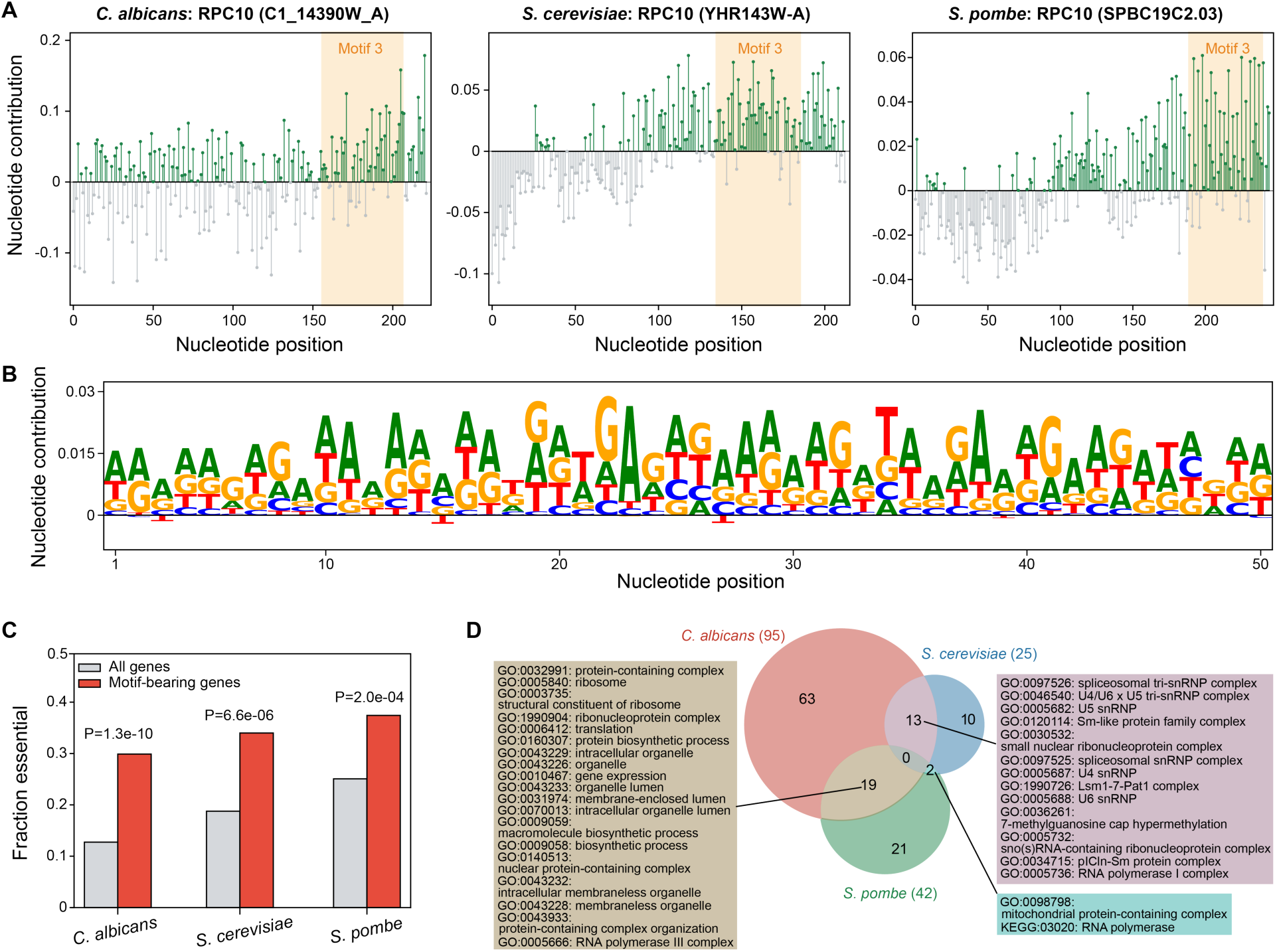
Nucleotide-level attribution and functional enrichment analysis of motifs associated with fungal gene essentiality. **A,** Nucleotide-level attribution scores across orthologous RPC10 coding sequences from *C. albicans*, *S. cerevisiae*, and *S. pombe*. Each stick represents the contribution of one nucleotide to the model’s predicted essentiality: green indicates a positive contribution and gray indicates a negative contribution. The orange-shaded region marks the position of pooled Motif 3. **B,** Attribution logo for pooled Motif 3, showing the mean signed nucleotide contribution pattern across aligned seqlets assigned to this motif. **C,** Fraction of essential genes among all genes (gray bars) and motif-bearing genes (red bars) in each species. P values were calculated using two-sided Fisher’s exact tests comparing the number of essential and non-essential genes between motif-bearing and non-motif-bearing gene sets within each species. **D,** Overlap of significantly enriched GO and KEGG terms among motif-bearing genes across the three species.

Despite limited motif conservation across species, motif-bearing genes were consistently enriched for essential genes in all three species (**Fig. 4C**). Functional enrichment analysis using Gene Ontology (GO) and Kyoto Encyclopedia of Genes and Genomes (KEGG) annotations further showed that these genes preferentially encoded core gene expression machinery, although no individual enriched term was shared across all three species (**Fig. 4D**; **Table S7**). In *C. albicans*, motif-bearing genes were enriched for translation- and ribosome-related functions, as well as spliceosomal small nuclear ribonucleoprotein (snRNP) complexes. In *S. cerevisiae*, enrichments were concentrated in spliceosomal and snRNP-related complexes, including U4/U6, U5, and tri-snRNP components. In *S. pombe*, motif-bearing genes were enriched primarily for ribosome- and ribonucleoprotein-complex-related GO/KEGG terms. Together, we found that gene essentiality in fungi is associated with coding sequence motifs enriched in genes encoding core gene expression and translation machinery, although these motifs are only partly conserved across species and remain predominantly species biased.

### Cross-species transfer learning enables gene essentiality prediction with little or no labeled data from the target species

Although our models performed well in *C. albicans*, *S. cerevisiae*, and *S. pombe*, training and testing within the same species does not address the challenge of predicting gene essentiality in a new species with little or no labeled data. To evaluate cross-species generalization, we trained models on two species and evaluated them on the third species (we refer these models as transfer learning models). Without using any essentiality labels from the target species, these models achieved mean AUROCs of 0.828-0.889 and mean AUPRCs of 0.570-0.698 (**Table S8**). We then used the transferred parameters to initialize each model and fine-tuned it with increasing number of labeled genes from the target species. Model performance improved as target species labels were added, but the incremental gains diminished rapidly with increasing training set size (**Fig. 5**, red curves). Thus, the pretrained models are already highly informative, and relatively few target species labels were required to achieve the best performance. In contrast, randomly initialized models performed substantially worse without target species labels and approached the performance of the transfer learning models only when larger amounts of labeled target species data were provided (**Fig. 5**, gray curves).

**Figure 5.**
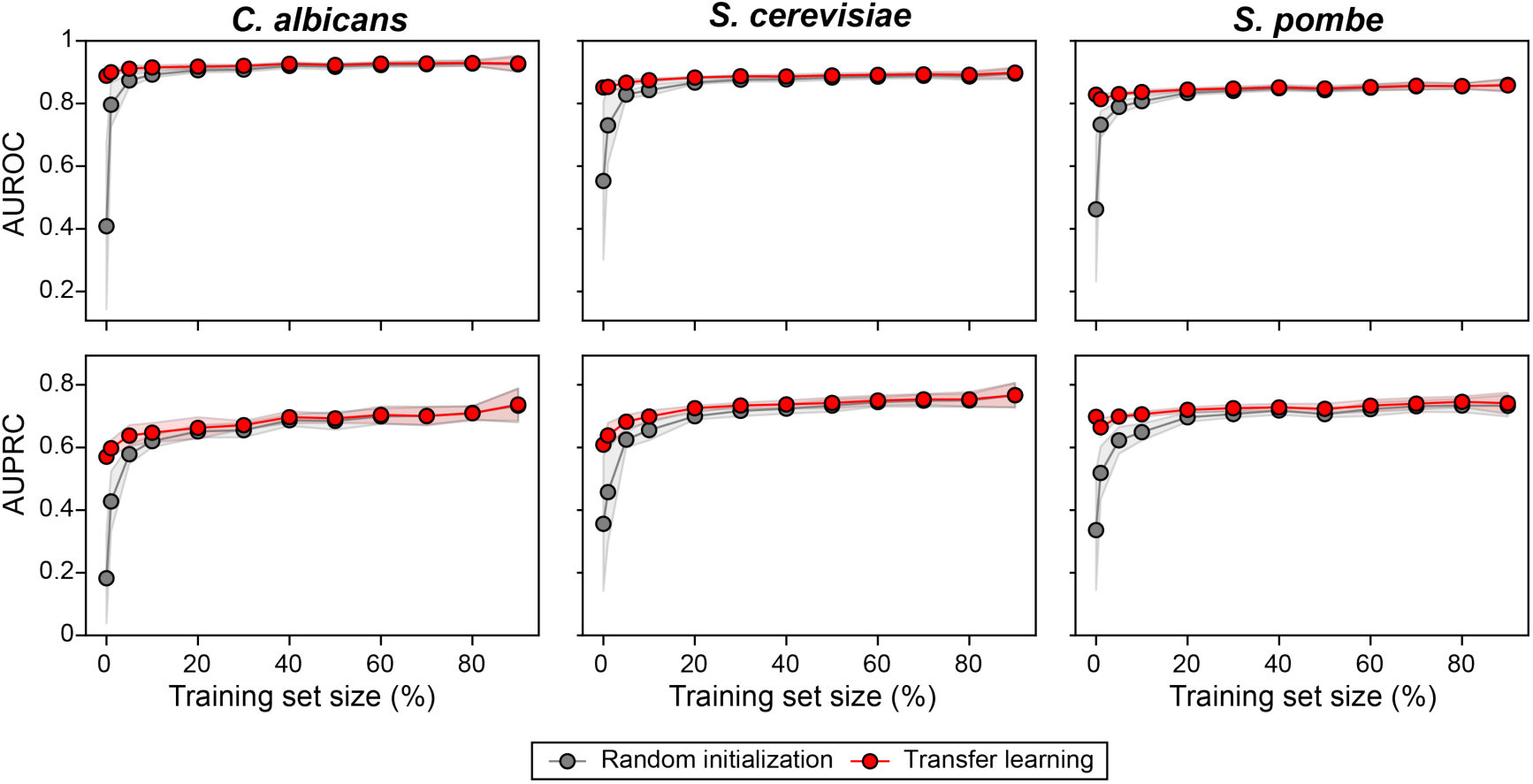
Cross-species gene essentiality prediction with increasing amounts of labeled target species data. Red dots and lines represent models initialized through transfer learning, whereas gray dots and lines represent models initialized randomly. Each dot represents the mean performance at a given target species training set size (expressed as a percentage of the total dataset) across 10 random splits. Shaded regions indicate ±1 standard deviation across the 10 splits.

We then extended this generalization test to an even more challenging prediction task: identifying changes in gene essentiality between strains of the same species. For this challenge, we selected 17 genes whose essentiality labels differ between *S. cerevisiae* strains S288C and W303 (**Fig. S3**; **see Methods**). Among the 11 genes that changed from essential in S288C to non-essential in W303, the predicted essentiality score decreased on average, although the fold change was modest and not significantly different from 1.0 (P = 0.378; *t*-test). In contrast, among the 6 genes that changed from non-essential in S288C to essential in W303, the predicted essentiality score increased by an average of 1.4-fold and was significantly greater than 1.0 (P = 0.044; *t*-test). Despite the challenge, our models captured the correct average direction of strain-specific essentiality changes, although not for every individual gene. Together, these results show that cross-species and -strain transfer learning improves gene essentiality prediction, even when target species training data are absent or very limited.

### Prediction of essential genes in *A. fumigatus* A1163

Finally, we applied our modeling framework to predict essential genes in *A. fumigatus* A1163, a major fungal pathogen causing life threatening infections in immunocompromised patients (28). We initialized the model using parameters learned from *C. albicans*, *S. cerevisiae*, and *S. pombe* and fine-tuned it using a small set of experimentally characterized genes. The positive set contains 35 genes identified as essential by conditional promoter replacement (11). The negative set includes 556 genes with viable gene replacement and null mutant strains maintained in the laboratory of Dr. Robert A. Cramer at Dartmouth’s Geisel School of Medicine. This dataset represents 5.97% of the 9,907 protein-coding genes in the A1163 genome, with essential genes comprising 5.92% of labeled genes.

Because the A1163 dataset is highly imbalanced, we tested whether this imbalance during fine-tuning affected model performance using a similar cross-species transfer learning framework used above (**see Methods**). The major difference was the class composition of the fine-tuning data: models were trained either on balanced data that preserve the essential-to-non-essential ratio of the full dataset or on imbalanced data matching the A1163 ratio. The resulting AUROC and AUPRC were similar between the two settings, indicating that class imbalance had little effect on ranking essential vs. non-essential genes. However, it substantially affected threshold-based classification. At the default essentiality score cutoff of 0.5, the imbalanced model had lower pooled recall (0.224 vs. 0.514) and F1 score (0.354 vs 0.600) when all species were combined. After optimizing the cutoff on held-out data, the imbalanced model required a lower threshold of 0.10 and achieved recall and F1 scores comparable to those of the balanced model (recall: 0.641 vs 0.639; F1 score: 0.631 vs 0.633).

Because the predicted scores were not well calibrated as probabilities, we used them to rank all 9,907 protein-coding genes and selected the top 1,500 highest-ranked genes as essentiality candidates (**Table S9**). This set represents 15.14% of the genome and includes genes with scores of at least 0.0901. Selecting 1,500 candidates also enabled direct comparison with a previous *A. fumigatus* prediction study by Lu *et al.*, which evaluated the top 1,500 predicted genes in the Af293 reference genome (23). Notably, the score threshold corresponding to the top 1,500 genes (0.0901) was very close to the optimal 0.10 threshold obtained in the above analyses of *C. albicans*, *S. cerevisiae*, and *S. pombe*. In addition, the resulting candidate proportion (15.14%) fell within the range of experimentally defined essential gene proportions in these three species (12.74%, 18.78%, and 25.07%, respectively).

To evaluate the candidate set, we compared it with candidate essential loci identified in an *A. fumigatus* Af293 parasexual transposon mutagenesis screen by Carr *et al.* (43). In that study, diploid transposon mutants were induced to generate haploid progeny, which were then tested for recovery of the transposon-bearing genome on selective medium. Loci for which no growing transposon-bearing haploids were recovered therefore provide evidence for essentiality, although the corresponding genes were not individually validated as strictly essential. From the 96 reported insertion loci, we reconstructed 90 unique protein-coding Af293 genes, among which 88 were mapped to A1163 with high confidence (**see Methods**). Of these, 52 (59.1%) genes were among our 1,500 candidates, including 3 used for fine-tuning (**Table S9**). This recovery was comparable to that reported by Lu *et al.*, in which a model trained on *Neurospora crassa* data recovered 50 of 90 genes (55.6%) among its own top 1,500 predictions.

To complement this screen-based evaluation, we conducted a separate literature-based assessment of model predictions (**see Methods**). We identified five experimentally verified essential genes from the A1163 lineage that were not included in the training or validation datasets: *pgm* (AFUB_037350) (44), *agm1* (AFUB_006590) (45), *pmmA* (AFUB_072510) (46), *uap1* (AFUB_088730) (47), and *gel4* (AFUB_022370) (48). Evidence for essentiality was obtained through conditional promoter replacement for *pgm*, *agm1*, *pmmA*, and *uap1,* and through heterokaryon rescue for *pmmA* and *gel4*. Four genes (*pgm*, *agm1*, *pmmA*, and *uap1*), which all encode enzymes in conserved pathways that generate nucleotide-sugar precursors for cell wall biosynthesis, scored above the 0.0901 threshold and were included in the 1500 predicted essential genes. In contrast, *gel4*, which encodes a β-1,3-glucanosyltransferase involved in cell wall remodeling during mycelial growth, received a score of 0.031 and was not included in the candidate set. Together, these results show that the model can recover experimentally validated essential genes involved in conserved cellular processes, while also revealing limitations in predicting essential genes associated with filamentous growth (see **Discussion**).

## Discussion

Our study demonstrates that GLM representations can support fungal gene essentiality prediction directly from DNA sequence. Evo2 embeddings captured essentiality-related signals even without task-specific training, while supervised training substantially improved performance over zero-shot scoring. Importantly, our modeling framework generalized across *C. albicans*, *S. cerevisiae*, and *S. pombe*, including transfer settings with limited or no target species data used for fine-tuning. This cross-species generalizability is the framework’s main practical value: for fungal species with available genomic sequences but no genome-wide essentiality data, transfer learning can provide an initial ranked list of candidate essential genes for experimental follow-up. We demonstrated this practical utility in *A. fumigatus*.

Our results also suggest that predictive performance was constrained more by the biological information contained in the input DNA representation than by the downstream classifier model capacity or type. Increasing MLP size or replacing the classifier head with alternative model families did not yield meaningful improvements (**Table 2**), whereas incorporating curated, biologically complementary features substantially improved performance (**Fig. 3**). Evo2 embeddings accurately captured sequence-proximal properties but were less effective at recovering features that reflect more distal, downstream biological processes or broader cellular context. Our findings thus suggest that, for biological sequence-to-function models, increasing model complexity alone may offer limited benefit when the input representation lacks information relevant to the function being predicted.

Although we did not identify a single universal motif shared across all three species, motif-bearing genes were enriched for functions involving core gene expression machinery. In *C. albicans*, several motifs displayed codon-position asymmetry in nucleotide contributions (**Fig. 4B**; **Fig. S4**), with stronger positive contributions at the first and third codon positions and weaker or negative contributions at the second position (**Fig. S5**). This triplet periodicity is consistent with a coding frame-dependent signal, suggesting that the model used features embedded in coding sequence architecture. Biologically, this pattern is plausible because the second codon position is often a major determinant of amino acid chemical properties and is therefore tightly constrained by protein function, whereas the third position is more flexible because it often tolerates synonymous substitutions. Third codon positions may thus encode signals related to codon usage, translational efficiency, mRNA stability, or other sequence features that distinguish essential genes. Notably, the periodicity is weaker or less apparent in *S. cerevisiae* and *S. pombe*, suggesting that the codon-level sequence features associated with essentiality differ among these species.

Application of our framework to *A. fumigatus* highlighted both the potential and limitations of cross-species transfer. The high rankings of *pgm*, *agm1*, *pmmA*, and *uap1* suggest successful transfer of conserved signals associated with carbohydrate metabolism and nucleotide-sugar production. In contrast, the low ranking of *gel4*, an essential β-1,3-glucanosyltransferase involved in mycelial cell wall remodeling, may reflect the difficulty of predicting essentiality associated with polarized hyphal growth and the distinct cell wall demands of mycelian development. This limitation is expected because, although *C. albicans* can form hyphae, the three reference species are primarily unicellular yeasts and may not capture the obligately filamentous, mycelial growth program of *A. fumigatus*. *N. crassa* is a closer taxonomic and morphological reference. However, the *N. crassa* knockout project classified genes as “probably essential” when deletion-bearing homokaryotic ascospores could not be recovered after sexual crosses (49). Because this endpoint cannot distinguish loss of viability from defects in sexual development, ascospore formation, or germination, we did not use these data for model training.

Several limitations of our study highlight important directions for future work. The DNA embeddings used here were derived from Evo2, a general GLM pretrained across all domains of life, including bacteria, archaea, and eukaryotes. We anticipate that fungal-specific GLMs may better capture fungal coding structure, genome composition, and regulatory context.

Although the contribution of PPI-derived features varied across species, the modular design of our framework allows updated or more accurate PPI prediction models to be incorporated as they become available. Future models could also integrate additional genomic sequence-derived predictions, including gene expression, protein localization, and regulatory network properties. Finally, gene essentiality is context dependent. Generating essentiality datasets under clinically relevant stresses, temperatures, and host-like conditions will be important for developing models that predict not only whether a gene is essential, but when and under what biological contexts it becomes essential.

## Materials and Methods

### Fungal essentiality datasets curation

Gene essentiality was treated as an assay- and condition-specific phenotype. We converted reported outcomes to a common gene-level classification: essential (1), non-essential (-1), or unresolved (0). Unknown, condition-dependent, and internally inconsistent outcomes were classified as unresolved and excluded from supervised model training and evaluation.

*C. albicans* essentiality data were obtained from Fu *et al.* (50), who used the Gene Replacement and Conditional Expression (GRACE) system. In this system, one allele of each gene is deleted and the remaining allele is placed under a repressible promoter, allowing gene requirement to be assessed from growth following transcriptional repression. For single-species modeling, we used phenotypes measured at 30°C. For cross-species analyses, we generated temperature-consensus labels by combining phenotypes measured at 30°C and 39°C. At each temperature, GRACE verdicts of “NE” (non-essential) and “GD” (growth defect) were treated as non-essential, whereas “E” was treated as essential. Concordant essential or non-essential calls were used to generate the consensus labels. When one temperature produced an unknown phenotype, the resolved phenotype from the other temperature was used. Genes with unknown phenotypes at both temperatures remained unknown. Discordant essential and non-essential calls were classified as condition-dependent, and any gene with an inconsistent call at either temperature was classified as unknown.

*S. cerevisiae* S288C essentiality data were obtained from the Saccharomyces Genome Database (SGD; https://www.yeastgenome.org/). SGD curates phenotypes from both standardized genome-wide mutant collections and gene-specific classical genetic studies, with experiment type terms describing the scale of the assay and, when specified, the ploidy of the mutant strain. The downloaded data were filtered to null mutants (Mutant_Type = null) and viability-related phenotypes. Because individual genes could have multiple annotations, we selected a single annotation using the following hierarchy, from highest to lowest priority: “systematic mutation set”, “homozygous diploid”, “systematic mutation set”, “large-scale survey”, “classical genetics”, “homozygous diploid”, and “heterozygous diploid”. For each gene, the highest-priority annotation was used as the essentiality label. “Inviable” mutants were classified as essential, whereas “viable”, “viability: normal”, “viability: decreased”, and “viability: increased” mutants were classified as non-essential because the null mutant remained viable under the reported assay conditions. The resulting dataset was filtered to include only genes present in the SGD S288C reference genome annotation.

*S. pombe* essentiality data were downloaded from PomBase (https://www.pombase.org/). The underlying deletion collection evaluates whether haploid spores carrying a deletion can germinate and establish viable colonies following sporulation of heterozygous diploids. Viability phenotypes were converted to essentiality labels and classified as essential (“inviable”), non-essential (“viable”), or unresolved (“condition-dependent”, “unknown”).

We assembled an *A. fumigatus* A1163 fine-tuning dataset comprising 35 experimentally validated essential genes (11) and 556 genes represented by viable knockout strains maintained in the Cramer laboratory. The 35 essential genes were tested in the CEA17 strain background (an A1163 lineage) by conditionally replacing their endogenous promoters with the nitrogen-regulated *niiA* promoter. Viable knockouts strains were considered evidence that the deleted gene was not required under the conditions used for strain construction and propagation. Data from the *N. crassa* knockout project (49) were not used in transfer learning because “probable essentiality” was assigned when deletion-bearing homokaryotic ascospores could not be recovered after sexual crossing heterokaryotic transformants. This outcome may reflect essentiality but can also result from defects in sexual development, ascospore formation, or germination.

### Evo2 scoring of ORFs and their perturbed coding sequences

For each *C. albicans* ORF, we generated two *in silico* perturbations: a coding-disruption mutation and a complete ORF deletion. For the coding-disruption mutation, a 15-nucleotide sequence containing five consecutive premature stop codons (TAATAATAATAGTGA) was inserted immediately after the first 12 nucleotides near the 5′ end of the ORF (27). For the deletion mutation, the entire coding sequence was removed.

Perturbations were evaluated using three sequence contexts: ORF-only, ORF plus available upstream and downstream intergenic sequences, and an ∼8 kb genomic context window capped at 8,192 nucleotides to match the context length of Evo2-7B-base. ORF plus intergenic sequences for *C. albicans* SC5314 were obtained from the Candida Genome Database Assembly 22 (https://www.candidagenome.org). To construct the ∼8 kb sequence context, flanking genomic sequence was added on both sides with the ORF centered whenever possible. For ORFs longer than 8,192 nucleotides, no flanking sequence was added. Instead, they were truncated at the 3’ end to retain the first 8,192 nucleotides.

Coding-disruption mutants were evaluated in all three settings. ORF-deletion mutants were evaluated only in the ORF-plus-intergenic and ∼8 kb genomic context settings because deleting the ORF leaves no sequence for ORF-only Evo2 scoring. For ORF deletions when an ∼8 kb genomic context was provided, the ORF was removed but its surrounding flanking sequence was retained. For each gene, perturbation, and context setting, Evo2-7B-base scores were computed for the wild-type and mutant sequences. Mutation effects were quantified as the mutant sequence score minus the corresponding wild-type sequence score. More negative values indicate a greater drop in model likelihood after mutation. We interpreted this drop as evidence of higher mutation intolerance and gene essentiality.

### Optimal Evo2 embedding generation

DNA sequences were represented using Evo2-7B-base embeddings. The optimal embedding generation configuration identified in **Fig. 2B** is reiterated here. For each gene, the ORF was embedded within an ∼8 kb genomic context window to match the 8,192-nucleotide context length used by Evo2-7B-base. Nucleotide-level embeddings were extracted from layer 26 of Evo2 and summarized into a single gene-level representation by mean pooling over ORF nucleotide positions only. This procedure produced a 4,096-dimensional DNA embedding vector for each gene.

### Ortholog essentiality features

Ortholog essentiality features were generated using ortholog relationships inferred by OrthoFinder v3.1.2 (38). For each target gene, we constructed an *N*-dimensional feature vector, where *N* is the number of reference species with available genome-scale essentiality datasets. Each element of the vector represents the essentiality status of the mapped orthologs in one reference species. For each reference species, the feature value was assigned as follows: 1 if any mapped ortholog was essential, -1 if at least one mapped ortholog was labeled non-essential and none was labeled essential, and 0 if no mapped ortholog had a binary essentiality label. During joint training across the three reference species, the feature corresponding to a gene’s own species was set to 0 to prevent label leakage.

### PPI network embeddings

Predicted PPI networks were generated from amino acid sequences using FlashPPI (36), specifically the tattabio/flashppi model, with a custom inference script based on the FlashPPI prediction workflow. Interactions with contact scores ≥ 0.5 were retained and used as edge weights to construct undirected PPI networks, in which proteins were represented as nodes. Network embeddings were then generated using weighted Node2Vec (39) with an embedding dimension of 128, walk length of 80, 10 walks per node, p = 1, q = 1, a random seed of 42, and one worker. The resulting random walks were used to train a skip-gram Word2Vec model with a window size of 5, min_count = 0, and one training epoch. Genes without a corresponding network embedding were assigned a 128-dimensional zero vector.

### Evo2-only and multimodal classifier architectures

The Evo2-only classifier is a MLP that receives frozen 4,096-dimensional Evo2 DNA embeddings as input. The network consists of layer normalization, a 4,096-to-256 linear layer, ReLU activation, and dropout (*p* = 0.3), followed by a 256-to-16 linear layer, ReLU activation, and dropout (*p* = 0.3). A final 16-to-1 linear layer produces a single essentiality logit.

The multimodal classifier integrates Evo2-derived DNA embeddings, predicted PPI-network embeddings, and ortholog-based essentiality features. DNA and PPI features are each encoded into 16-dimensional latent representations by separate two-layer MLP encoders. Each encoder consists of layer normalization, a linear projection from the input dimension (4,096 for DNA or 128 for PPI) to 256 dimensions, ReLU activation, dropout (*p* = 0.3), a 256-to-16 linear layer, and ReLU activation. The *N*-dimensional ortholog feature vector was projected directly to a 16-dimensional representation with a linear layer.

Pairwise interactions between the DNA and PPI, DNA and ortholog, and PPI and ortholog representations were modeled using three independent bilinear layers. Each bilinear layer generates a 16-dimensional interaction vector followed by tanh activation. The three modality-specific latent vectors and three pairwise interaction vectors were concatenated to form a 96-dimensional representation, which was passed through a prediction head comprising a 96-to-16 linear layer, dropout (*p* = 0.3), and a final 16-to-1 linear layer that produces an essentiality logit.

### Evo2-only and multimodal classifier training and evaluation

Unless otherwise specified, Evo2-only and multimodal essentiality classifiers were evaluated using the same stratified five-fold cross validation splits. Within each training fold, 20% of the genes were reserved by stratified sampling for validation and early stopping. Models were trained for up to 500 epochs using binary cross-entropy loss with logits, the AdamW optimizer, a learning rate of 4 × 10^-5^, weight decay of 0.05, a batch size of 256, and gradient clipping with a maximum norm of 1.0. Training was stopped when validation loss did not improve for 20 consecutive epochs, and the checkpoint with the lowest validation loss was used to evaluate the held-out test fold. Test set logits were converted to essentiality scores using the sigmoid function. AUROC and AUPRC were calculated separately for each test fold and reported as the mean across the five folds.

### Cross-species and strain-level transfer learning

To evaluate cross-species transfer learning, we trained the multimodal model on labeled genes from two species of *C. albicans*, *S. cerevisiae*, and *S. pombe* and transferred the learned parameters to the third, held-out species. The transferred model was evaluated either directly, without target species labels, or after fine-tuning with 1%-90% of the labeled target species genes. At each training set size, performance on the remaining labeled genes was evaluated across 10 stratified, class-balanced random splits and compared with models initialized with random weights and trained using the same target species data. To evaluate transfer between

*S. cerevisiae* strains, we used 80% of the labeled *S. cerevisiae* S288C genes for model training and the remaining 20% for validation and early stopping. The trained model was then applied to W303 directly. Comparison of SGD-derived essentiality labels identified 19 genes with different classifications between the two strains. Two genes, *YBR135W* and *YLR099W-A*, were absent from the W303 reference peptide FASTA file and were therefore excluded, leaving 17 genes. For each gene, the predicted strain-specific change was calculated as the ratio of its essentiality score in W303 relative to that in S288C. The score ratios were analyzed separately for genes changing in each direction (non-essential to essential, and essential to non-essential) and compared with 1.0 (baseline, meaning no change) using a one-sample *t* test.

### Classifier capacity and model family benchmarking

MLP classifier capacity was evaluated by varying the numbers of neurons in its two hidden layers while keeping all other architectural and training settings unchanged. The first and second hidden layers contained 64 and 8 neurons, 256 and 16 neurons, 1,024 and 64 neurons, or 4,096 and 256 neurons, respectively. The MLP was also compared with logistic regression, k-nearest neighbors, random forest, and XGBoost classifiers, which were trained and evaluated using the same data splits. Logistic regression was fit with a maximum of 1,000 iterations. Default random forest and XGBoost models were fit using 1,000 estimators. For optimized random forest and XGBoost models, hyperparameters were tuned separately within each training fold by inner three-fold stratified cross validation, and the parameter combination with the highest mean average precision was selected.

### Essentiality prediction using curated features and leave-one-feature-out analysis

Using values for 13 curated features obtained from Fu *et al.* (14), we trained an MLP to predict *C. albicans* gene essentiality. The model consists of input batch normalization, a 64-neuron hidden layer with ReLU activation and dropout (*p* = 0.3), and a sigmoid output layer that produces an essentiality score. Models were evaluated using the same five predefined cross validation splits used for *C. albicans* essentiality prediction. Within each training fold, 20% of the genes were randomly reserved for validation and early stopping. Models were trained for up to 500 epochs using binary cross-entropy loss, the Adam optimizer, a learning rate of 1 × 10^-4^, weight decay of 0.05, and a batch size of 256. Training was stopped after 20 consecutive epochs without improvement in validation loss, and the checkpoint with the lowest validation loss was used to evaluate the held-out test fold. Leave-one-feature-out analysis was performed by removing each curated feature individually and repeating the same five-fold evaluation. AUROC and AUPRC were reported as the mean of the five fold-specific values.

### Recovery of 13 curated features from Evo2 embeddings

Values of the 13 curated features were obtained from Fu *et al.* (14). We trained a separate MLP regression probe to predict each feature from Evo2 embeddings. Models were evaluated using the same five predefined cross validation splits used for *C. albicans* essentiality prediction. Within each fold, feature values were z-standardized using the training set, and the same transformation was applied to the held-out test set. 20% of the training set was randomly reserved for validation and early stopping. Each probe used the same architecture as the Evo2-only MLP classifier, except that the final layer produced a continuous value rather than an essentiality logit. Probes were trained for up to 500 epochs using mean squared error loss, the AdamW optimizer, a learning rate of 4 × 10^-5^, weight decay of 0.05, a batch size of 256, and gradient clipping with a maximum norm of 1.0. Training was stopped after 20 consecutive epochs without improvement in validation loss, and the checkpoint with the lowest validation loss was used to evaluate the held-out test fold. Spearman’s correlation coefficient (ρ) was calculated separately between predicted and observed standardized values in each held-out test fold, and feature recoverability was reported as the mean of the five fold-specific ρ values.

### Input modality ablation and fusion strategy analysis

We evaluated the contribution of each input modality and their combinations by systematically removing one or more modalities from the multimodal model. The full model incorporated Evo2 DNA embeddings, ortholog-based essentiality, and PPI-network features. In the “no PPI”, “no DNA”, and “no ortholog” models, the corresponding encoder and all bilinear interaction terms involving that modality were removed. In single-modality models (“DNA only”, “ortholog only”, and “PPI only”), the prediction head was applied solely to the latent representation of the retained modality. To compare fusion strategies, we retained all three input modalities and varied only the fusion module. In the concatenation model, the three 16-dimensional modality-specific latent representations were directly concatenated into a 48-dimensional vector without explicit interaction terms. In the cross-attention model, the three latent representations were stacked and processed using scaled dot-product attention with learned query, key, and value projections. Self-attention was masked so that each modality attended only to the other two modalities. The resulting context vectors were transformed with tanh, concatenated with the original modality-specific representations, and passed to the prediction head.

### Nucleotide-level attribution by integrated gradients

Integrated gradients (41) were computed using Captum v0.8.0 (51) with 50 integration steps. The Evo2 DNA embedding tensor for each gene was provided as the model input, whereas ortholog-based essentiality features and PPI-network embeddings were supplied as fixed additional arguments. Captum generated signed attribution values for each embedding dimension at each nucleotide position. These values were summed across embedding dimensions to yield a single signed contribution score per nucleotide position, which was used as the importance profile for motif discovery. Zero-valued embedding tensors matching the shape of the DNA embedding input were used as baselines.

### Motif discovery and functional enrichment analysis

Motif discovery was performed using a TF-MoDISco-based workflow (42) implemented in modiscolite v2.4.0 (https://github.com/jmschrei/tfmodisco-lite). We first ran motif discovery separately for *C. albicans*, *S. cerevisiae*, and *S. pombe*. For each species, one-hot-encoded DNA sequences were combined with signed nucleotide-level attribution scores to generate contribution tracks. Positive high-importance seqlets were identified using identical conservative parameters across species, including a target seqlet FDR of 0.005, a minimum passing-window fraction of 0.0005, a maximum passing-window fraction of 0.002, a minimum metacluster size of 40, and a final minimum cluster size of 15. Seqlets were then clustered independently within each species into recurrent DNA motifs. To assess whether shared motifs could be recovered across species, we also pooled the input sequences and attribution scores from all three species and reran the same TF-MoDISco workflow using the same seqlet-discovery parameters. For each pooled motif, we calculated the proportion of its seqlets originating from each species to determine whether the pooled motif was disproportionately represented by one species. To determine whether pooled motifs were associated with conserved gene families, we assigned motif-bearing genes to OrthoFinder orthogroups. For each motif, we then identified orthogroups containing at least one motif-bearing gene from each of the three species. Motif 3 was the only pooled motif associated with an orthogroup meeting this criterion.

To test whether motif-bearing genes were enriched for essential genes, we compared the fraction of essential genes among motif-bearing genes with the genome-wide background using two-sided Fisher’s exact tests. Functional enrichment of motif-bearing genes was performed with g:Profiler (52) using GO biological process, molecular function, cellular component, and KEGG annotations. Enriched GO/KEGG terms were identified separately for each species and then compared by term identifier to determine species-specific and shared functional categories.

### Evaluation of class imbalance effects

For each target species (*C. albicans*, *S. cerevisiae*, or *S. pombe*), we initialized the model with parameters learned from the other two species, following the cross-species transfer-learning procedure used in **Fig. 5**. We generated 10 random fine-tuning splits. Within each split, the balanced and imbalanced training conditions were evaluated on the same held-out test set comprising 50% of the labeled genes. The number of genes in the training sets was limited to 5.97% of all labeled genes, matching the proportion of labeled genes available for *A. fumigatus* A1163. Under the balanced condition, training genes were sampled to preserve the species-specific essential-to-non-essential ratio. Under the imbalanced condition, training genes were sampled to match the assembled *A. fumigatus* A1163 dataset label distribution (5.92% essential vs. 94.08% non-essential). Within each training set, 20% of genes were reserved for validation during model training. The optimal essentiality score cutoff was defined as the threshold that maximized the F1 score on the held-out test set.

### *A. fumigatus* A1163 model training and evaluation

The *A. fumigatus* A1163 model was initialized with parameters from a multimodal model trained on combined data of *C. albicans*, *S. cerevisiae*, and *S. pombe*. The pretrained model on these three species used a single stratified 80% training/20% validation split of the pooled labeled genes, with the validation set used for early stopping. No separate test set was used at this stage. This model was subsequently fine-tuned using the assembled *A. fumigatus* dataset. This dataset includes 35 essential genes from Hu *et al.* (11) and 556 non-essential genes with viable knockouts maintained in the Cramer laboratory. Among these 556 non-essential genes, four gene identifiers (AFUB_004190, AFUB_042770, AFUB_049550, and AFUB_069160) could not be matched to the 9,907 protein-coding genes in FungiDB release 68 (3). Three genes were absent from the annotation, and one was annotated as a pseudogene. These genes were thus excluded from fine-tuning. The resulting dataset of 587 genes was divided into 80% training and 20% validation sets using stratified sampling.

We compared our A1163 essentiality rankings with the candidate essential loci reported by Carr *et al.* (43) in the CEA17-derived diploid strains CEA225 and CEA226. Carr *et al.* reported protein-coding genes associated with these insertion sites using *Afua_* locus identifiers, which correspond to the Af293 reference annotation. Their Table 2 contains 96 insertion records, with four protein-coding genes represented by two independent insertions each. After collapsing these duplicate gene assignments, we excluded the signal recognition particle (SRP) RNA locus and the record listing *Afua_1g05240* (the locus affected by the insertion was unannotated and *Afua_1g05240* was the nearest annotated gene). This yielded 90 unique Af293 protein-coding gene annotations associated with the Carr *et al.* insertion sites. Candidate A1163 orthologs (*AFUB_* identifiers) were obtained from the FungiDB release 68 Orthologs table. We independently compared the complete Af293 and A1163 proteomes by bidirectional BLASTP (53), retaining only matches with E-values ≤ 1 × 10^-10^. Ortholog assignments were considered high confidence when the Af293 and A1163 proteins were reciprocal best hits (defined as the highest bit score match in both search directions) and when the same pair was designated syntenic in FungiDB. Both criteria were met by 88 of the 90 annotations. Recovery was defined as the number of these 88 mapped genes occurring among the 1,500 highest-ranked A1163 genes.

### Literature search for *A. fumigatus* essential genes

To further evaluate our fine-tuned *A. fumigatus* A1163 model, we performed a literature search for experimentally validated essential genes. Searches were conducted in PubMed, Web of Science, and ASM Journals using the following terms: (“A1163” OR “Aspergillus fumigatus” OR “A. fumigatus”) AND (“essential” OR “critical for viability”) AND (“Tet-OFF” OR “null mutant” OR “heterokaryon rescue”). Studies were included if they used strains in the A1163 lineage (CEA10, CEA17, A1163, ΔakuBKU80, or A1160) and provided direct experimental evidence for essentiality. Eligible validation approaches included creation of a conditional mutant such as Tet-OFF and heterokaryon rescue. Studied were excluded if they used other strain backgrounds (e.g., AfS77, A1280, ATCC46645, Af237), inferred essentiality solely from transposon-mutagenesis datasets, or examined genes included in the model training or validation datasets.

### Use of generative AI in manuscript preparation

During manuscript preparation, the authors used OpenAI ChatGPT and Codex to assist with language editing and improve the clarity and readability of author-generated text. All AI-assisted output was reviewed and revised by the authors, who take full responsibility for the final content.

## Data availability

All data used to generate the tables and figures are included in the manuscript and supplementary materials. Customized Python code for training the machine learning models, along with the final pretrained model and fine-tuning scripts for new fungal species or genomes, is available on GitHub at https://github.com/HannahGT-289/Gene_Essentiality.git.

## Acknowledgements

This study was funded by NIH/NIAID grant DP2AI200956 (C.L.), NIH/NIAID grant R00AI175599 (C.L.), NIDDK P30 grant DK117469 (Dartmouth Cystic Fibrosis Research Center), and startup funds from the Department of Microbiology and Immunology at Dartmouth College (C.L.).

All authors declare no conflict of interest.

C.L. initiated and designed the project. C.L. and H.G.T. developed, implemented, and evaluated the machine learning models, analyzed the data, and interpreted the results. All authors drafted, reviewed, edited, and approved the final manuscript.

## References

1. Wang X-W, Wang T, Liu Y-Y. 2026. Artificial intelligence for microbiology and microbiome research. Cell Syst 17:101531.

2. Feldbauer R, Schulz F, Horn M, Rattei T. 2015. Prediction of microbial phenotypes based on comparative genomics. BMC Bioinform 16:S1.

3. Basenko EY, Shanmugasundram A, Böhme U, Starns D, Wilkinson PA, Davison HR, Crouch K, Maslen G, Harb OS, Amos B, McDowell MA, Kissinger JC, Roos DS, Jones A. 2024. What is new in FungiDB: a web-based bioinformatics platform for omics-scale data analysis for fungal and oomycete species. Genetics 227:iyae035.

4. Nambiar A, Dubinkina V, Liu S, Maslov S. 2023. FUN-PROSE: A deep learning approach to predict condition-specific gene expression in fungi. PLOS Comput Biol 19:e1011563.

5. Khaiwal S, De Chiara M, Barré BP, Barrio-Hernandez I, Stenberg S, Beltrao P, Warringer J, Liti G. 2025. Predicting natural variation in the yeast phenotypic landscape with machine learning. Mol Syst Biol 21:1466–1489.

6. Shahreen N, Osinuga A, Malla S, Razmpour T, Tabibian M, Saha R. 2026. Multi-omics integration in genome-scale metabolic models: a review of constraint-based approaches. Mol Omics 22:aaiag005.

7. Rancati G, Moffat J, Typas A, Pavelka N. 2018. Emerging and evolving concepts in gene essentiality. Nat Rev Genet 19:34–49.

8. Batté A, Bosch-Guiteras N, Pons C, Ota M, Lopes M, Sharma S, Tellini N, Paltenghi C, Conti M, Kan KT, Ho UL, Wiederkehr M, Barraud J, Ashe M, Aloy P, Liti G, Chabes A, Parts L, Leeuwen J van. 2026. The modifiers that cause changes in gene essentiality. Cell Syst 17:101515.

9. Hillenmeyer ME, Fung E, Wildenhain J, Pierce SE, Hoon S, Lee W, Proctor M, St.Onge RP, Tyers M, Koller D, Altman RB, Davis RW, Nislow C, Giaever G. 2008. The chemical genomic portrait of yeast: uncovering a phenotype for all genes. Science 320:362–365.

10. Costanzo M, VanderSluis B, Koch EN, Baryshnikova A, Pons C, Tan G, Wang W, Usaj M, Hanchard J, Lee SD, Pelechano V, Styles EB, Billmann M, van Leeuwen J, van Dyk N, Lin Z-Y, Kuzmin E, Nelson J, Piotrowski JS, Srikumar T, Bahr S, Chen Y, Deshpande R, Kurat CF, Li SC, Li Z, Usaj MM, Okada H, Pascoe N, San Luis B-J, Sharifpoor S, Shuteriqi E, Simpkins SW, Snider J, Suresh HG, Tan Y, Zhu H, Malod-Dognin N, Janjic V, Przulj N, Troyanskaya OG, Stagljar I, Xia T, Ohya Y, Gingras A-C, Raught B, Boutros M, Steinmetz LM, Moore CL, Rosebrock AP, Caudy AA, Myers CL, Andrews B, Boone C. 2016. A global genetic interaction network maps a wiring diagram of cellular function. Science 353:aaf1420.

11. Hu W, Sillaots S, Lemieux S, Davison J, Kauffman S, Breton A, Linteau A, Xin C, Bowman J, Becker J, Jiang B, Roemer T. 2007. Essential Gene Identification and Drug Target Prioritization in *Aspergillus fumigatus*. PLoS Pathog 3:e24.

12. Becker JM, Kauffman SJ, Hauser M, Huang L, Lin M, Sillaots S, Jiang B, Xu D, Roemer T. 2010. Pathway analysis of *Candida albicans* survival and virulence determinants in a murine infection model. Proc Natl Acad Sci U S A 107:22044–22049.

13. Segal ES, Gritsenko V, Levitan A, Yadav B, Dror N, Steenwyk JL, Silberberg Y, Mielich K, Rokas A, Gow NAR, Kunze R, Sharan R, Berman J. 2018. Gene Essentiality Analyzed by In Vivo Transposon Mutagenesis and Machine Learning in a Stable Haploid Isolate of *Candida albicans*. mBio 9:e02048–18.

14. Fu C, Zhang X, Veri AO, Iyer KR, Lash E, Xue A, Yan H, Revie NM, Wong C, Lin Z-Y, Polvi EJ, Liston SD, VanderSluis B, Hou J, Yashiroda Y, Gingras A-C, Boone C, O’Meara TR, O’Meara MJ, Noble S, Robbins N, Myers CL, Cowen LE. 2021. Leveraging machine learning essentiality predictions and chemogenomic interactions to identify antifungal targets. Nat Commun 12:6497.

15. Billmyre RB, Craig CJ, Lyon JW, Reichardt C, Kuhn AM, Eickbush MT, Zanders SE. 2025. Landscape of essential growth and fluconazole-resistance genes in the human fungal pathogen *Cryptococcus neoformans*. PLOS Biol 23:e3003184.

16. Seringhaus M, Paccanaro A, Borneman A, Snyder M, Gerstein M. 2006. Predicting essential genes in fungal genomes. Genome Res 16:1126–1135.

17. Campos TL, Korhonen PK, Gasser RB, Young ND. 2019. An Evaluation of Machine Learning Approaches for the Prediction of Essential Genes in Eukaryotes Using Protein Sequence-Derived Features. Comput Struct Biotechnol J 17:785–796.

18. Nigatu D, Sobetzko P, Yousef M, Henkel W. 2017. Sequence-based information-theoretic features for gene essentiality prediction. BMC Bioinform 18:473.

19. Gustafson AM, Snitkin ES, Parker SC, DeLisi C, Kasif S. 2006. Towards the identification of essential genes using targeted genome sequencing and comparative analysis. BMC Genom 7:265.

20. Lu H, Li F, Yuan L, Domenzain I, Yu R, Wang H, Li G, Chen Y, Ji B, Kerkhoven EJ, Nielsen J. 2021. Yeast metabolic innovations emerged via expanded metabolic network and gene positive selection. Mol Syst Biol 17:e10427.

21. Cheng J, Wu W, Zhang Y, Li X, Jiang X, Wei G, Tao S. 2013. A new computational strategy for predicting essential genes. BMC Genom 14:910.

22. Beder T, Aromolaran O, Dönitz J, Tapanelli S, Adedeji EO, Adebiyi E, Bucher G, Koenig R. 2021. Identifying essential genes across eukaryotes by machine learning. NAR Genom Bioinform 3:lqab110.

23. Lu Y, Deng J, Rhodes JC, Lu H, Lu LJ. 2014. Predicting essential genes for identifying potential drug targets in *Aspergillus fumigatus*. Comput Biol Chem 50:29–40.

24. Schonfeld E, Vendrow E, Vendrow J, Schonfeld E. 2021. On the relation of gene essentiality to intron structure: a computational and deep learning approach. Life Sci Alliance 4.

25. Benegas G, Ye C, Albors C, Li JC, Song YS. 2025. Genomic language models: opportunities and challenges. Trends Genet 41:286–302.

26. Consens ME, Dufault C, Wainberg M, Forster D, Karimzadeh M, Goodarzi H, Theis FJ, Moses A, Wang B. 2025. Transformers and genome language models. Nat Mach Intell 7:346– 362.

27. Brixi G, Durrant MG, Ku J, Naghipourfar M, Poli M, Sun G, Brockman G, Chang D, Fanton A, Gonzalez GA, King SH, Li DB, Merchant AT, Nguyen E, Ricci-Tam C, Romero DW, Schmok JC, Taghibakhshi A, Vorontsov A, Yang B, Deng M, Gorton L, Nguyen N, Wang NK, Pearce MT, Simon E, Adams E, Amador ZJ, Ashley EA, Baccus SA, Dai H, Dillmann S, Ermon S, Guo D, Herschl MH, Ilango R, Janik K, Lu AX, Mehta R, Mofrad MRK, Ng MY, Pannu J, Ré C, St. John J, Sullivan J, Tey J, Viggiano B, Zhu K, Zynda G, Balsam D, Collison P, Costa AB, Hernandez-Boussard T, Ho E, Liu M-Y, McGrath T, Powell K, Pinglay S, Burke DP, Goodarzi H, Hsu PD, Hie BL. 2026. Genome modelling and design across all domains of life with Evo 2. Nature 652:1349–1361.

28. James MR, Doss KE, Cramer RA. 2024. New developments in *Aspergillus fumigatus* and host reactive oxygen species responses. Curr Opin Microbiol 80:102521.

29. Firon A, Villalba F, Beffa R, d’Enfert C. 2003. Identification of Essential Genes in the Human Fungal Pathogen *Aspergillus fumigatus* by Transposon Mutagenesis. Eukaryot Cell 2:247–255.

30. Romero B, Turner G, Olivas I, Laborda F, Ramón De Lucas J. 2003. The *Aspergillus nidulans alcA* promoter drives tightly regulated conditional gene expression in *Aspergillus fumigatus* permitting validation of essential genes in this human pathogen. Fungal Genet Biol 40:103–114.

31. Firon A, Beauvais A, Latgé J-P, Couvé E, Grosjean-Cournoyer M-C, d’Enfert C. 2002. Characterization of essential genes by parasexual genetics in the human fungal pathogen *Aspergillus fumigatus*: impact of genomic rearrangements associated with electroporation of DNA. Genetics 161:1077–1087.

32. Kang B, Fan R, Cui C, Cui Q. 2025. Comprehensive prediction and analysis of human protein essentiality based on a pretrained large language model. Nat Comput Sci 5:196–206.

33. Ma J, Song J, Young ND, Chang BCH, Korhonen PK, Campos TL, Liu H, Gasser RB. 2024. ‘Bingo’—a large language model- and graph neural network-based workflow for the prediction of essential genes from protein data. Brief Bioinform 25:bbad472.

34. Zotenko E, Mestre J, O’Leary DP, Przytycka TM. 2008. Why Do Hubs in the Yeast Protein Interaction Network Tend To Be Essential: Reexamining the Connection between the Network Topology and Essentiality. PLOS Comput Biol 4:e1000140.

35. Taha K. 2025. Protein-protein interaction detection using deep learning: A survey, comparative analysis, and experimental evaluation. Comput Biol Med 185:109449.

36. Cornman A, Tranzillo M, Zulaybar NG, Bouzit I, Hwang Y. 2026. Linear-time prediction of proteome-scale microbial protein interactions. Proc Natl Acad Sci U S A 123:e2610619123.

37. Szklarczyk D, Nastou K, Koutrouli M, Kirsch R, Mehryary F, Hachilif R, Hu D, Peluso ME, Huang Q, Fang T, Doncheva NT, Pyysalo S, Bork P, Jensen LJ, von Mering C. 2025. The STRING database in 2025: protein networks with directionality of regulation. Nucleic Acids Res 53:D730–D737.

38. Emms DM, Liu Y, Belcher L, Holmes J, Kelly S. 2026. OrthoFinder: improved phylogenetic orthology inference with enhanced accuracy and scalability. Nat Methods 23:1327–1333.

39. Grover A, Leskovec J. 2016. node2vec: Scalable Feature Learning for Networks, p. 855–864. *In* Proceedings of the 22nd ACM SIGKDD International Conference on Knowledge Discovery and Data Mining. Association for Computing Machinery, New York, NY, USA.

40. Tsai Y-HH, Bai S, Liang PP, Kolter JZ, Morency L-P, Salakhutdinov R. 2019. Multimodal Transformer for Unaligned Multimodal Language Sequences, p. 6558–6569. In Korhonen, A, Traum, D, Màrquez, L (eds.), Proceedings of the 57th Annual Meeting of the Association for Computational Linguistics. Association for Computational Linguistics, Florence, Italy.

41. Sundararajan M, Taly A, Yan Q. 2017. Axiomatic Attribution for Deep Networks, p. 3319–3328. *In* Proceedings of the 34th International Conference on Machine Learning. PMLR.

42. Shrikumar A, Tian K, Avsec Ž, Shcherbina A, Banerjee A, Sharmin M, Nair S, Kundaje A. 2018. Technical Note on Transcription Factor Motif Discovery from Importance Scores (TF-MoDISco) version 0.5.6.5. arXiv.org. https://arxiv.org/abs/1811.00416v5. Retrieved 2 August 2026.

43. Carr PD, Tuckwell D, Hey PM, Simon L, d’Enfert C, Birch M, Oliver JD, Bromley MJ. 2010. The Transposon *impala* Is Activated by Low Temperatures: Use of a Controlled Transposition System To Identify Genes Critical for Viability of *Aspergillus fumigatus*. Eukaryot Cell 9:438–448.

44. Yan K, Stanley M, Kowalski B, Raimi OG, Ferenbach AT, Wei P, Fang W, Aalten DMF van. 2022. Genetic validation of *Aspergillus fumigatus* phosphoglucomutase as a viable therapeutic target in invasive aspergillosis. J Biol Chem 298.

45. Fang W, Du T, Raimi OG, Hurtado-Guerrero R, Mariño K, Ibrahim AFM, Albarbarawi O, Ferguson MAJ, Jin C, Van Aalten DMF. 2013. Genetic and structural validation of *Aspergillus fumigatus* N-acetylphosphoglucosamine mutase as an antifungal target. Biosci Rep 33:e00063.

46. Zhang Y, Fang W, Raimi OG, Lockhart DEA, Ferenbach AT, Lu L, van Aalten DMF. 2021. Genetic and structural validation of phosphomannomutase as a cell wall target in *Aspergillus fumigatus*. Mol Microbiol 116:245–259.

47. Fang W, Du T, Raimi OG, Hurtado-Guerrero R, Urbaniak MD, Ibrahim AFM, Ferguson MAJ, Jin C, van Aalten DMF. 2013. Genetic and structural validation of *Aspergillus fumigatus* UDP-N-acetylglucosamine pyrophosphorylase as an antifungal target. Mol Microbiol 89:479– 493.

48. Gastebois A, Fontaine T, Latgé J-P, Mouyna I. 2010. β(1-3)Glucanosyltransferase Gel4p Is Essential for *Aspergillus fumigatus*. Eukaryot Cell 9:1294–1298.

49. Colot HV, Park G, Turner GE, Ringelberg C, Crew CM, Litvinkova L, Weiss RL, Borkovich KA, Dunlap JC. 2006. A high-throughput gene knockout procedure for *Neurospora* reveals functions for multiple transcription factors. Proc Natl Acad Sci U S A 103:10352–10357.

50. Fu C, Xiong EH, Kupczok L, Archambault LS, Wang TRW, Holleran C, Carruthers-Lay D, Zhuang TX, Marcoccia S, Zhang H, Chen K, Anderson D, Yiu B, Liu Z, Herzel L, Robbins N, Cowen LE. 2025. Expansion of the functional genomics GRACE library reveals genes relevant for temperature-dependent fitness in *Candida albicans*. PLOS Biol 23:e3003409.

51. Kokhlikyan N, Miglani V, Martin M, Wang E, Alsallakh B, Reynolds J, Melnikov A, Kliushkina N, Araya C, Yan S, Reblitz-Richardson O. 2020. Captum: A unified and generic model interpretability library for PyTorch. arXiv:2009.07896. arXiv 10.48550/arXiv.2009.07896.

52. Kolberg L, Raudvere U, Kuzmin I, Adler P, Vilo J, Peterson H. 2023. g:Profiler— interoperable web service for functional enrichment analysis and gene identifier mapping (2023 update). Nucleic Acids Res 51:W207–W212.

53. Camacho C, Coulouris G, Avagyan V, Ma N, Papadopoulos J, Bealer K, Madden TL. 2009. BLAST+: architecture and applications. BMC Bioinform 10:421.

